# Differences between apoplastic and cytosolic reactive oxygen species production in *Arabidopsis* during pattern-triggered immunity

**DOI:** 10.1101/2021.04.20.440614

**Authors:** Dominique Arnaud, Michael J. Deeks, Nicholas Smirnoff

**Affiliations:** Biosciences, College of Life and Environmental Sciences, University of Exeter, Exeter EX4 4QD, UK; Laboratory of Molecular Plant Physiology and Functional Genomics and Proteomics of Plants, CEITEC-Central European Institute of Technology, Masaryk University, Kamenice 5, CZ-625 00, Brno, Czech Republic.

**Keywords:** Pathogen associated molecular patterns, oxidative burst, stomata, redox sensors

## Abstract

Despite an ever-increasing interest in reactive oxygen species (ROS) signalling during plant-microbe interactions, very little information exists, mainly for technical reasons, on the molecular mechanisms regulating intracellular hydrogen peroxide (H_2_O_2_) signalling during PAMP-triggered immunity. Here, we used a sensitive fluorimetry method and the H_2_O_2_ sensor roGFP2-Orp1, which revealed unsuspected features on the regulation of cytoplasmic H_2_O_2_ and thiol redox dynamics upon pathogen-associated molecular patterns (PAMPs) perception by *Arabidopsis thaliana*. Extended PAMP-induced cytosolic roGFP2-Orp1 oxidation was distinct from the transient oxidative burst in the apoplast measured by luminol oxidation. Pharmacological and genetic analyses indicate that the prolonged PAMP-induced H_2_O_2_ increase in the cytoplasm was largely independent on NADPH oxidases and apoplastic peroxidases. By contrast, the NADPH oxidase mutant *rbohF* was hyper-sensitive to roGFP2-Orp1 oxidation by H_2_O_2_ and PAMP indicating a lower antioxidant capacity. Unlike previous reports, the *rbohF* mutant, but not *rbohD*, was impaired in PAMP-triggered stomatal closure and ROS production measured by a fluorescein-based probe in guard cells resulting in defects in stomatal defences against bacteria. However, stomatal closure was not correlated with an increase in roGFP2-Orp1 oxidation in guard cells. Interestingly, RBOHF also participated in PAMP-induced apoplastic alkalinisation. Altogether, our results provide novel insights on the interplay between apoplastic and cytosolic ROS dynamics and highlight the importance of RBOHF in plant immunity.

**Significance statement:** Plants mount defence responses to pathogens by detecting pathogen-associated molecular patterns (PAMPs). One response is a rapid and transient burst of reactive oxygen species (ROS, e.g. superoxide and hydrogen peroxide) in the cell wall (apoplast) produced by NADPH oxidases and cell wall peroxidases. Using a genetically-encoded hydrogen peroxide sensor roGFP2-Orp1, we found that, in contrast to the transient apoplastic ROS burst, there is also prolonged hydrogen peroxide production in the cytosol upon PAMP perception which is independent of NADPH oxidase and cell wall peroxidases. Our results suggest that apoplastic ROS rather than intracellular hydrogen peroxide is a signal triggering stomatal closure during PAMP-triggered immunity. Additionally, we re-address the relative contribution of the NADPH oxidases D and F in stomatal immunity.

## Introduction

Reactive oxygen species (ROS) such as hydrogen peroxide (H_2_O_2_) are signalling molecules involved in various biological processes such as development and responses to environmental stresses. The production of ROS in response to pathogens and pathogen-associated molecular patterns (PAMPs) is common across many groups of organisms. In plants, an early PAMP-triggered immunity (PTI) response within minutes, is a transient apoplastic oxidative burst mediated by plasma membrane NADPH oxidases (termed RBOH in plants) and also cell wall peroxidases (PRXs) (1–4). Extracellularly produced H_2_O_2_ diffuses into the cell probably through aquaporin to activate downstream defence responses such as stomatal closure or callose deposition (5, 6). The perception of PAMPs by plasma membrane receptor kinases (RKs) activates the co-receptor BAK1 and the cytosolic kinase BIK1 which in turn phosphorylates and activates RBOHD (7, 8). NADPH oxidase uses cytoplasmic NADPH as an electron source, transported *via* FAD and heme cofactors to the outside where oxygen is reduced to superoxide. The majority of this superoxide is assumed to dismutate very rapidly, producing hydrogen peroxide (9). While RBOHD is the main NADPH oxidase isoform involved in apoplastic ROS burst and participates in stomatal (pre-invasive) defence responses, its contribution to plant resistance against *Pseudomonas syringae* pv tomato (*Pst*) bacteria is not clear (4, 7, 10). On the other hand, RBOHF partly involved in apoplastic ROS production (3, 4) is required for full post-invasive (late apoplastic) resistance against virulent *Pst* bacteria (10). Type III cell wall peroxidases, while using hydrogen peroxide to oxidatively cross-link cell wall components, can under pathogen perception also generate ROS/H_2_O_2_ (1, 2). In particular, PRX4, PRX33, PRX34 and PRX71 are involved in PAMP-mediated ROS production, and PRX33/34 play an important role in pre- and post-invasive defences against *Pst* bacteria (2, 11). Although, the molecular mechanisms activating PRXs upon PAMP perception are not yet resolved, apoplastic alkalinisation is a prerequisite for PRX-mediated ROS production (12).

The chemical probes generally used in measuring PAMP and pathogen-associated ROS production include luminol, diaminobenzidine (DAB) and 2’,7’ dichlorofluorescein diacetate (H_2_DCFDA). These mostly lack spatial resolution and, more importantly, specificity as they can react with a range ROS such as superoxide ions (O_2_^•-^), peroxynitrite (ONOO^-^), and hydroxyl radical (OH^•^) (9). To improve detection of H_2_O_2_, genetically encoded biosensors such as HyPer or roGFP2-Orp1 have been characterized *in vitro* and *in vivo* using diverse model organisms and are being increasingly used in plants (13–18). The main advantages of these sensors are the possibility to make ratiometric measurements that are independent of the level of probe expression, and the reversibility of the probe oxidation by the glutaredoxin (GRX) and glutathione (GSH) mediated reduction permitting dynamic and real-time measurements. Furthermore, given this interaction with the thiol system, probes reactive with H_2_O_2_ can be compared with those that report the redox state of the glutathione pool (19).

Recently, *Arabidopsis thaliana* expressing roGFP2-Orp1 in the cytosol/nucleus or mitochondria has been used to study H_2_O_2_ accumulation in leaves exposed to PAMPs in mutants affected in GSH/GRX redox metabolism (18), showing that this probe could be successfully used to monitor the plant immune response. Here, we investigated the role of NADPH oxidases, PRXs, and upstream PTI regulators BAK1 and BIK1 on intracellular changes in H_2_O_2_/redox dynamics during PAMP-triggered immunity by crossing mutants with Arabidopsis plants expressing the cytosolic/nuclear roGFP2-Orp1 biosensor. Whole leaf responses were measured by fluorimetry and stomatal guard cell-specific responses by confocal laser scanning microscopy. Unexpectedly, our results reveal that PAMP-mediated H_2_O_2_ accumulation in the cytosol/nucleus of leaves is largely independent on apoplastic ROS produced by NADPH oxidases and PRXs. Despite an increase in H_2_O_2_ level in guard cells upon PAMP perception, roGFP2-Orp1 oxidation state in guard cells was not tightly correlated with stomatal aperture. Importantly, we found that RBOHF plays a crucial role in stomatal immunity by controlling ROS production and apoplastic pH.

## Results

### Intracellular H_2_O_2_ dynamics is influenced by the NADPH oxidase RBOHF and apoplastic peroxidases PRX4, PRX33 and PRX34

The response of the H_2_O_2_ biosensor roGFP2-Orp1 to diverse oxidant and reductant species has been thoroughly characterised *in vitro* and *in vivo* in *Arabidopsis thaliana* (14, 18, 20). To add to this information, we determined the *in vivo* H_2_O_2_ dose response of roGFP2-Orp1, localised to the cytoplasm/nucleus (18), using the multi-well fluorimetry method with leaf discs from rosette leaves of mature Arabidopsis plants. The roGFP2-Orp1 fluorescence emission at 505-545 nm was measured after sequential dual excitation at 400 nm and 485 nm and the ratio 400/485 nm was expressed relative to the initial ratio (Ri) measured before treatment. Dose response experiments indicated that the cytosolic/nuclear roGFP2-Orp1 is immediately oxidized by exogenous H_2_O_2_ in a dose-dependent manner between 100 μM and 100 mM (Fig. 1A and B) and 10 μM H_2_O_2_ was insufficient to oxidise the probe. As observed before (18, 21), roGFP2-Orp1 was reduced by DTT but the small decrease from the control treatment R/Ri ratio (Fig. 1B) suggests that the probe is largely reduced in untreated leaf discs and indicates its suitability to investigate changes in H_2_O_2_ levels upon treatment by different stimuli.

**Fig. 1.**
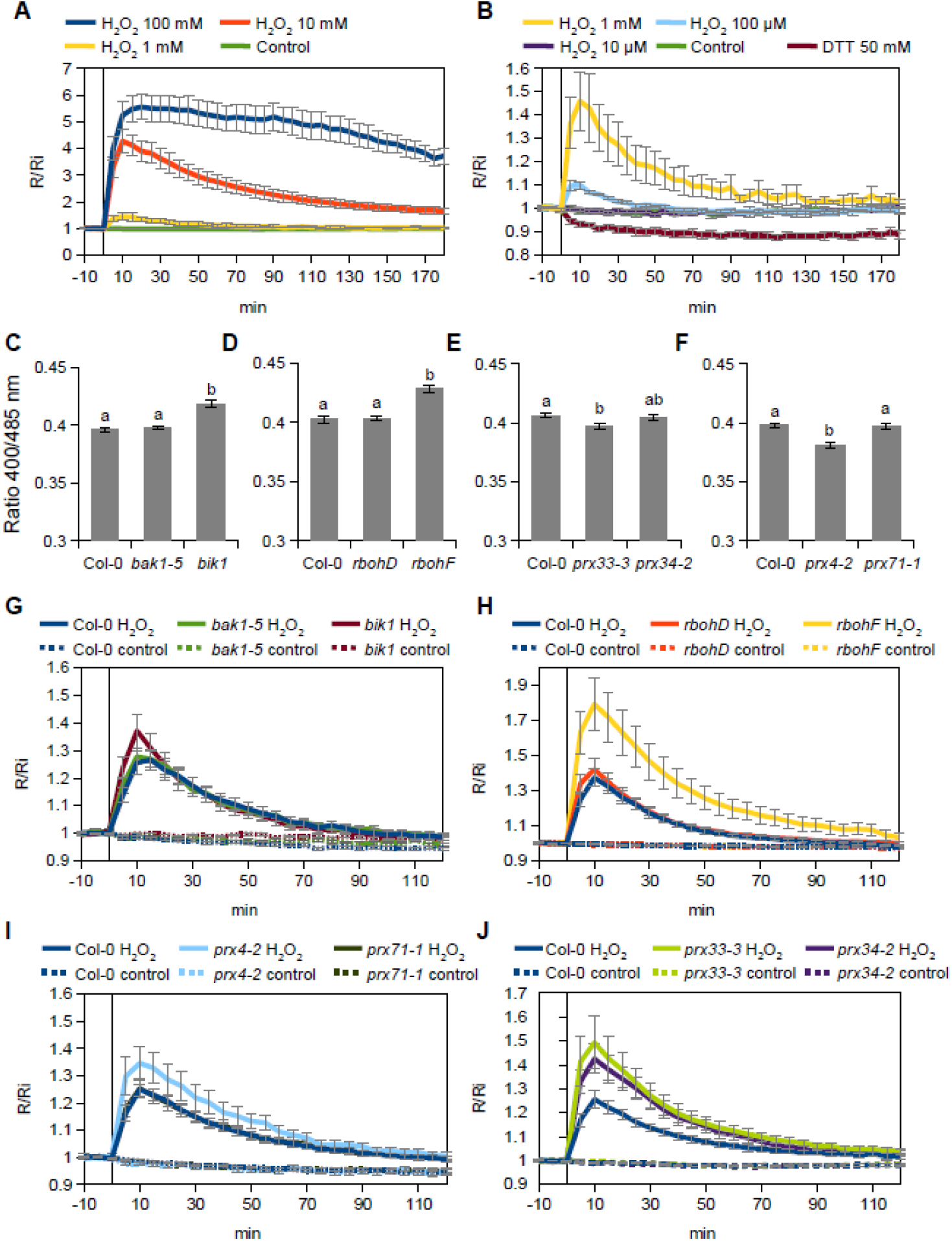
Initial oxidation state and H_2_O_2_-induced oxidation of roGFP2-Orp1 in mutants of PTI regulators, NADPH oxidases and apoplastic peroxidases. (A-B) *In vivo* characterisation of roGFP2-Orp1 oxidation and reduction kinetics in response to H_2_O_2_ and DTT. Leaf discs from rosette leaves of 5 week-old plants were exposed at t = 0 min to control solution, various concentrations of H_2_O_2_, or 50 mM DTT. The 400/485 nm fluorescence ratio (R) was measured over time by multiwell fluorimetry (excitation at 400 ± 8 and 485 ± 8 nm; emission, 525 ± 20 nm) and expressed relative to the mean initial ratio (Ri) before treatment (R/Ri). For clarity high concentrations of H_2_O_2_ are shown in (A) and lower doses are shown in (B). Data are means ± SE from a representative experiment (n = 6). The experiments have been repeated at least twice with similar results. (C-F) Initial roGFP2-Orp1 oxidation state in *bak1-5* and *bik1* (C), *rbohD* and *rbohF* (D), *prx33-3* and *prx34-2* (E) and *prx4-2* and *prx71-1* (F) mutants. The oxidation state of cytoplasmic/nuclear roGFP2-Orp1 (ratio 400/485 nm) in untreated condition was measured by multiwell fluorimetry on leaf discs. Data are means ± SE of at least three independent experiments (n ≥ 40). Different letters indicate significant differences at *P* < 0.001 (A and B), *P* < 0.05 (C) and *P* < 0.01 (D) based on a Tukey’s HSD test. (G-J) Kinetics of roGFP2-Orp1 oxidation in leaves of *bak1-5* and *bik1* (G), *rbohD* and *rbohF* (H), *prx33-3* and *prx34-2* (I) and *prx4-2* and *prx71-1* (J) mutants in response to exogenous H_2_O_2_. Leaf discs were exposed at t = 0 min to control solution or 1 mM H_2_O_2_, the 400/485 nm fluorescence ratio (R) was measured over time by multiwell fluorimetry and expressed relative to the mean initial ratio (Ri) before treatment. Data are means ± SE from 3 independent experiments (n ≥ 15).

To study how PAMP-induced extracellular ROS affects intracellular H_2_O_2_ levels and downstream physiological responses such as stomatal closure, Arabidopsis expressing cytosol/nucleus localised roGFP2-Orp1 (18) was crossed with mutants affected in upstream regulators of plant immunity BAK1 and BIK1 (22–24) and genes known to be involved in ROS production in the apoplast such as the NADPH oxidases RBOHD and RBOHF (3, 4, 10) and the apoplastic peroxidases PRX4, PRX33, PRX34 and PRX71 (2, 11). Note that contrary to the other lines used which are knock-out mutants, the *prx4-2* mutation in the 3’UTR induces overexpression of *PRX4* (11) and the *bak1-5* mutant is specifically impaired in PTI responses due to a mis-sense nucleotide substitution (25).

In untreated leaf discs, roGFP2-Orp1 was unexpectedly more oxidised in *rbohF* and *bik1* mutant backgrounds than in Col-0 wild-type (WT), while the oxidation state of the biosensor was not affected in *bak1-5* and *rbohD* mutants (Fig 1C and D). This suggests a lower antioxidant capacity in the *bik1* and *rbohF* mutants, possibly due to long term adaptation to the lack of O_2_^·-^ production by RBOHF. By contrast, roGFP2-Orp1 was significantly more reduced in *prx33-3* than WT suggesting that PRX33 produces H_2_O_2_ even in unstressed conditions (Fig. 1E). On the contrary, the stronger reduction of roGFP2-Orp1 in the *prx4-2* background (Fig. 1F) indicates that PRX4 scavenges H_2_O_2_ for the polymerisation of phenolic compounds (26). These differences in the extent of roGFP2-Orp1 oxidation, depending on the mutant backgrounds prompted an investigation of the effect of exogenous H_2_O_2_. Compared to WT, roGFP2-Orp1 became more oxidised in *bik1, prx4-2, prx33-3, prx34-2* and particularly *rbohF*, in response to 1 mM H_2_O_2_ (Fig. 1G-J). The hypersensitivity of *prx33-3* and *prx34-2* mutants to H_2_O_2_ could be explained by a defect in H_2_O_2_ removal, as these apoplastic peroxidases are also known to consume H_2_O_2_ to catalyse the oxidation of monolignols for lignin polymerisation (27).

### BAK1, BIK1, RBOHD, PRX4 and PRX34 influence PAMP-induced roGFP2-Orp1 oxidation

To investigate intracellular H_2_O_2_ dynamics during PAMP-triggered immunity, we analysed the responsiveness of roGFP2-Orp1 to PAMP treatments. A dose response assay indicated that roGFP2-Orp1 is oxidized by the PAMP flagellin 22 (flg22) at a concentration as low as 10 nM (Fig. 2A), which is comparable to other physiological assays using this elicitor (28). Probe oxidation started after a lag of 10 minutes and reached a plateau at 60 min, remaining oxidised over the 180 min duration of the experiment. GRX1-roGFP was similarly oxidised but with a somewhat larger change compared to roGFP-Orp1 (Fig. 2B). The PAMP elf18 peptide at 1 μM elicited a similar roGFP2-Orp1 oxidation response to flg22 (Fig. S1).

**Fig. 2.**
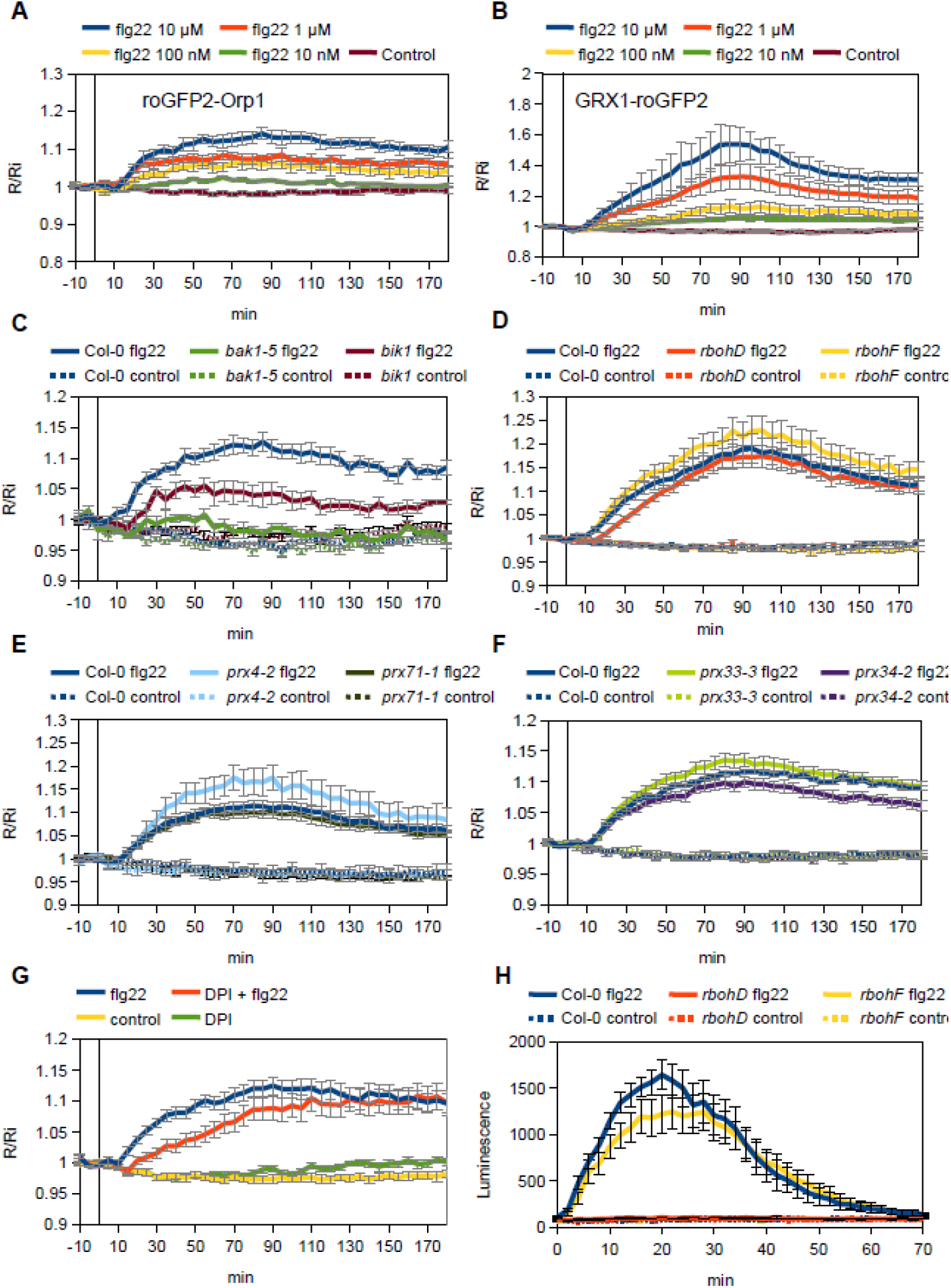
flg22-induced intracellular roGFP2-Orp1 redox dynamics in mutants affecting PTI-mediated ROS production in the apoplast. (A-B) Dose response kinetics of roGFP2-Orp1 (A) and GRX1-roGFP2 (B) oxidation in leaves in response to the PAMP flg22. Leaf discs were exposed at t = 0 min to control solution or different concentration of flg22. The 400/485 nm fluorescence ratio (R) was measured over time by multiwell fluorimetry and expressed relative to the mean initial ratio (Ri) before treatment. Data are means ± SE from a representative experiment (n = 6). The experiments have been repeated at least twice with similar results. (C-F) Kinetics of roGFP2-Orp1 oxidation in leaves of *bak1-5* and *bik1* (B), *rbohD* and *rbohF* (C), *prx33-3* and *prx34-2* (D) and *prx4-2* and *prx71-1* (E) mutants in response to flg22. Leaf discs were exposed at t = 0 min to control solution or 1 μM flg22, the ratio 400/485 nm (R) was measured over time by multiwell fluorimetry and expressed relative to the mean initial ratio (Ri) before treatment. Data are means ± SE from three independent experiments (n ≥ 15, C-E). In (B), a representative experiment is shown (n = 6). (G) Effect of DPI on flg22-induced oxidation of roGFP2-Orp1. After 2 hrs of pre-treatment with control solution or 20 μM DPI, leaf discs from Col-0 were exposed at t = 0 min to control solution or 1 μM flg22. The 400/485 nm fluorescence ratio (R) was measured over time by multiwell fluorimetry and expressed relative to the mean initial ratio (Ri) before flg22 treatment. Data are means ± SE of three independent experiments (n ≥ 10). (H) PAMP-induced apoplastic ROS production detected by luminol assay in Col-0 WT, *rbohD* and *rbohF* mutants. The luminescence was measured over time after treatment with control solution or 1 μM flg22 at t= 0 min. Data are means ± SE (n = 6) from a representative experiment.

The *bak1-5, bik1, rbohD, rbohF, prx4-2,prx33-3,prx34-2* and *prx71-1* mutants were subjected to flg22 or elf18 treatments (Fig. 2C-D and Fig. S1). The *bik1* and *bak1-5* mutants were strongly impaired in PAMP-triggered roGFP2-Orp1 oxidation (Fig. 2C and Fig. S1A). Notably, *rbohD* had a significantly delayed PAMP response, not reaching the oxidation level of Col-0 until 50 minutes after addition of either flg22 (Fig. 2D) or elf18 (Fig. S1B) respectively. Correspondingly, pre-treatment with the flavoenzyme/NADPH oxidase inhibitor diphenyleneiodonium chloride (DPI) mimicked the response of *rbohD* by causing a significant lag in flg22-induced roGFP2-Orp1 oxidation (Fig. 2G). roGFP-Orp1 oxidation kinetics were compared to the PAMP-induced apoplastic oxidative burst using a luminol assay (28). This oxidative burst showed the expected fast and transient response, peaking at 20 minutes post flg22 addition and was absent in *rbohD* (Fig. 2H). Therefore, the delayed roGFP-Orp1 oxidation in *rbohD* corresponds to the time of the “missing” apoplastic burst. By contrast, *rbohF, prx33-3* and *prx71-1* were not affected in PAMP-induced roGFP2-Orp1 oxidation while the *prx34-2* mutant exhibited a slight reduction of roGFP2-Orp1 oxidation from 100 min after PAMP treatments (Fig. 2D-F and Fig S1B-D). The *prx4-2* over-expression line showed an enhanced oxidation of roGFP2-Orp1 after PAMP (and H_2_O_2_) treatments (Fig. 2E and Fig S1C), indicating that PRX4 can produce H_2_O_2_ under stress conditions that over-accumulates in the cytosol.

roGFP2-Orp1 is also oxidised *in vitro* by peroxynitrite (ONOO^-^) and hypochlorous acid (HOCl), but not by nitric oxide (NO) (20), While HOCl production has not been reported in plants, peroxynitrite is formed from the reaction between NO and superoxide (O_2_^·-^) which are both produced during plant defence responses and programmed cell death (29, 30). Therefore, to assess the potential role of peroxynitrite in PAMP-induced roGFP2-Orp1 oxidation, NO supply was manipulated using the NO donor sodium nitroprusside (SNP) and the NO scavenger carboxyphenyl-4,4,5,5-tetramethylimidazoline-1-oxyl 3-oxide (cPTIO) (31). roGFP2-Orp1 was oxidised *in vivo* by high SNP concentrations (>5 mM, Fig S2A) but not by lower concentrations (< 0.5 mM) which is in the range used in typical experiments (32, 33). A similar response was observed with the *E_GSH_* biosensor GRX1-roGFP2 (Fig S2B) which was reported to be much less efficiently oxidised by ONOO^-^ than roGFP2-Orp1 *in vitro* (20). Surprisingly, two hours of pre-treatment with cPTIO induced oxidation of both roGFP2-Orp1 and GRX1-roGFP2 (Fig S3A and S3C). However, the flg22-induced roGFP2-Orp1 oxidation was not affected by cPTIO (Fig S3B), while GRX1-roGFP2 oxidation was moderately decreased by cPTIO at later time points (Fig S3D). These results suggest that the increase in roGFP2-Orp1 oxidation by flg22 reflects mainly the production of H_2_O_2_ rather than NO or ONOO^-^.

### The *rbohF* mutant is affected in flg22-mediated apoplastic alkalinisation

Apoplastic alkalinisation is a well characterised early event induced upon PAMP perception (28, 34) and constitutively active plasma membrane H^+^ATPase AHA1 impedes PAMP-mediated stomatal closure and apoplastic ROS burst (35, 36). Otte and collaborators showed that elicitor-induced alkalinisation of the apoplast was inhibited by DPI in chickpea (37) and NADPH oxidases could contribute to extracellular pH changes through their electrogenic activities and/or the consumption of protons during superoxide dismutation (38). To test this hypothesis, we used the ratiometric Oregon green fluorescent dye which has been successfully used in Arabidopsis (39, 40) and adapted the method for multi-well fluorimetry on leaf discs. Interestingly, assays with Oregon green indicate that the apoplastic pH was higher in the *rbohF* mutant in unstressed conditions while it remained unchanged in *rbohD* background (Fig. 3A). This result nicely correlates with the impaired growth of *rbohF* rosette leaves observed previously (3, 10) and in our laboratory conditions suggesting that RBOHF is important for the acidic growth of plant cells. Importantly, *rbohF* mutant was partly defective in PAMP-induced apoplastic alkalinisation but *rbohD* showed WT increase in pH after flg22 treatment (Fig. 3B).

**Fig. 3.**
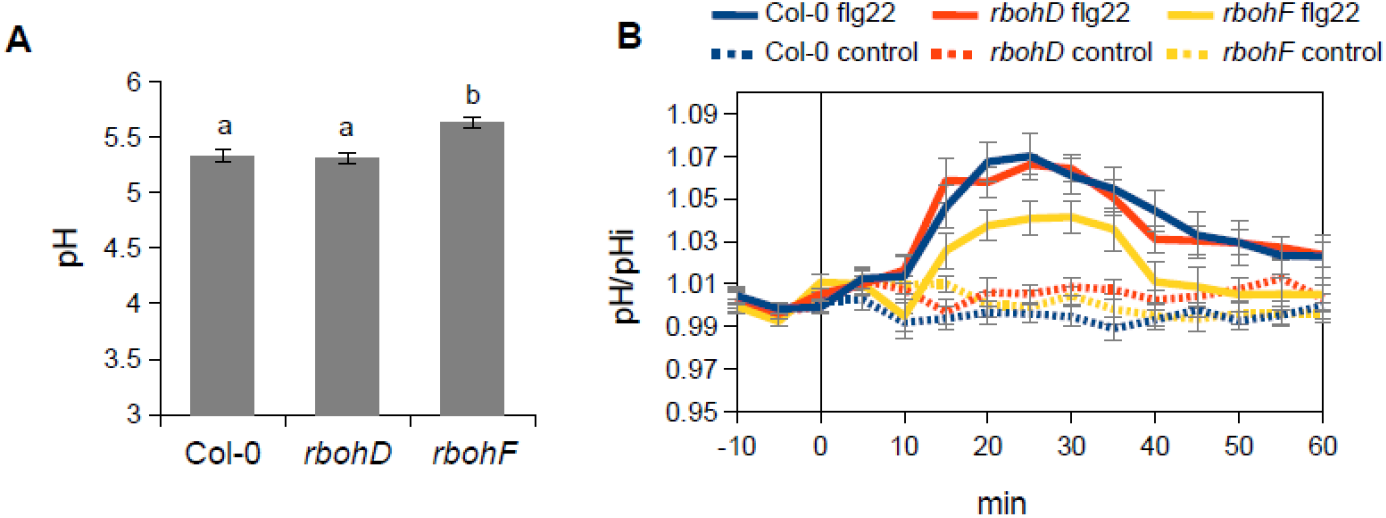
RBOHF influences apoplastic pH (A) Apoplastic pH in Col-0 WT, *rbohD* and *rbohF* mutant leaves. Apoplastic pH in untreated condition was determined by multiwell fluorimetry (sequential excitation at 440 ± 8 and 495 ± 8 nm; emission, 525 ± 20 nm) on Oregon green dextran-infiltrated leaf discs from 5 week-old plants (See Methods for details). Data are means ± SE (n ≥ 12) from a representative experiment. Different letters indicate significant differences at *P* < 0.001 based on a Tukey’s HSD test. (B) Kinetics of flg22-induced leaf apoplastic alkalinisation in Col-0 WT, *rbohD* and *rbohF* mutants. Oregon green dextran-infiltrated leaf discs were exposed at t = 0 min to control solution or 1 μM flg22, the apoplastic pH was measured over time by multiwell fluorimetry and expressed relative to the mean initial pH (pHi) before treatment (pH/pHi). Data are means ± SE (n ≥ 18) from 3 independent experiments.

### RBOHF is involved in stomatal defence responses

Stomatal closure is one of the first lines of defence against foliar bacteria (41) and PAMP-mediated stomatal closure has been correlated with increased ROS production in guard cells (42, 43). Both NADPH oxidase (RBOHD) and apoplastic peroxidases (PRX33 and PRX34) participate in stomatal immunity (11, 44, 45). Indeed, pre-treatment with an inhibitor of NADPH oxidases (DPI) or with salicylhydroxamic acid (SHAM) or sodium azide two inhibitors of apoplastic peroxidases (46) compromised flg22-mediated stomatal closure (Fig. S4A). Stomatal aperture was analysed in *rbohD* and *rbohF* leaf discs after treatment with flg22 or inoculation with the coronatine-deficient *Pseudomonas syringae* pv. tomato DC3000 (*Pst* COR^-^) bacteria for 2 hrs (Fig. 4A). A comparison with epidermal peels was also made (Fig. S4B-C). Because the toxin coronatine counteracts PAMP-mediated stomatal closure, the *Pst* COR^-^ strain has been widely used to characterise mutants defective in stomatal immunity (7, 41, 42, 45). Contrary to previous observations showing that RBOHD is required for PAMP-induced stomatal closure (7, 8, 44, 45), we found that in our experimental conditions *rbohD* consistently exhibited WT stomatal closure upon PAMP or bacterial treatments on both leaf disc and epidermal peel assays (Fig. 4A and Fig. S4B-C). On the contrary, the *rbohF* mutant was clearly defective in PAMP- and bacteria-mediated stomatal closure (Fig. 4A and Fig. S4B-C). These results were confirmed for flg22 by using different mutant alleles of these NADPH oxidases (Fig. 4B). Although *RBOHD* is reported to be more expressed in guard cells than *RBOHF* (47, 48), RT-qPCR measurements show that, contrary to *RBOHD, RBOHF* expression was induced in guard cells 2 hrs after flg22 treatment (Fig. S4D). Compared with the PAMP-mediated induction of *RBOHD* expression in young seedlings (47), our results indicate tissue specific induction of these two NADPH oxidases after PAMP activation and highlight the importance of RBOHF in stomatal immunity.

**Fig. 4.**
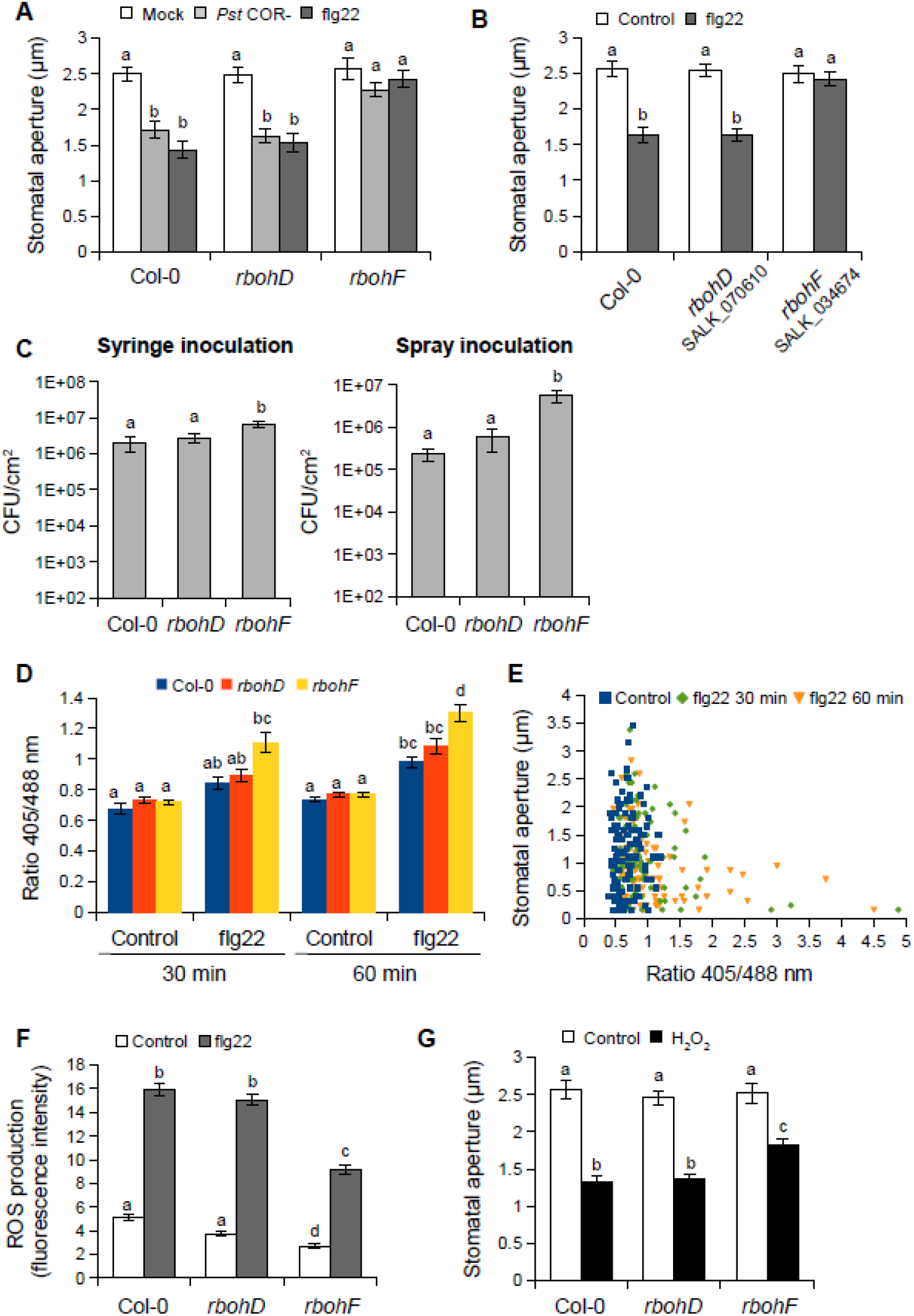
Analysis of the function of RBOHD and RBOHF during stomatal defence responses (A) Stomatal apertures in WT Col-0, *rbohD*, and *rbohF* leaf discs exposed to Mock control (10 mM MgCl2), 10^8^ cfu/ml COR-deficient *Pst* DC3000 (*Pst* COR^-^) bacteria and 5 μM flg22 for 2 h. Data are means ± SE (n ≥ 100) from a representative experiment. (B) Stomatal apertures in Col-0 WT and other allelic mutants of *rbohD* (SALK_070610) and *rbohF* (SALK_034674). Epidermal peels were exposed to Control solution or 5 μM flg22 for 2 h. Data are means ± SE (n ≥ 80) from a representative experiment. (C) Bacterial growth in WT Col-0, *rbohD*, and *rbohF* mutants assessed at 3 days after syringe-infiltration with *Pst* COR^-^ at 10^6^ cfu/ml (left panel) or after spray-inoculation with *Pst* COR^-^ at 10^8^ cfu/ml (right panel). Values are the means ± SE (n = 6). (D) Oxidation state of roGFP2-Orp1 in Col-0 WT, *rbohD* and *rbohF* guard cells in response flg22. Leaf discs were exposed to control solution or 1 μM flg22 and the ratio 405/488 nm of stomata was quantified at 30 min and 60 min after treatment from images of the fluorescence emission at 517 ± 9 nm following excitation at 488 and 405 nm. Data are means ± SE (n ≥ 50 guard cells) from a representative experiment. (E) Correlation between the stomatal aperture and the oxidation state of roGFP2-Orp1 in guard cells of leaf discs after treatment with control solution or 1 μM flg22 for 30 min and 60 min. The scatter plots show the stomatal aperture as a function of the ratio 405/488 nm for each condition (n ≥ 100). The Pearson correlation coefficient (R) is 0.0651 (*P* = 0.483) for control treatment and −0.2428 (*P* = 0.015) and −0.2395 (*P* = 0.011) for flg22 treatment at 30 min and 60 min respectively. (F) ROS production detected by H_2_DCFDA fluorescence in guard cells of Col-0 WT, *rbohD*, and *rbohF* epidermal peels 30 min after treatment with control solution or 5 μM flg22. Data are means ± SE (n ≥ 200) from 3 independent experiments. (G) Stomatal apertures in WT Col-0, *rbohD*, and *rbohF* leaf discs exposed to Control, or 100 μM H_2_O_2_ for 2 h. Data are means ± SE (n ≥ 60) from a representative experiment. Different letters indicate significant differences at *P* < 0.001 (A, B and G) and *P* < 0.05 (C, D and F) based on a Tukey’s HSD test.

Because a defect in stomatal defence often results in enhanced susceptibility to bacteria (Melotto et al., 2006), leaves were spray-inoculated with *Pst* COR^-^ bacteria and bacterial growth was evaluated 3 days later (Fig 4C). The *rbohF* mutant was more susceptible than Col-0 control to surface inoculated bacteria, with a 24-fold increase in bacterial population. As observed before (7, 45) the *rbohD* mutant was as susceptible as Col-0 to surface inoculated *Pst* COR^-^ corroborating the absence of defects in bacteria-mediated stomatal closure. To verify any deficiency in post-invasive defences, plants were also inoculated with *Pst* COR^-^ by syringe infiltration. Similar to previous results with the WT *Pst* DC3000 bacteria (10), the *rbohD* mutant exhibited WT susceptibility to bacteria while *rbohF* was slightly more susceptible to syringe-inoculated *Pst* COR^-^ with a 3-fold increase in bacterial titer as compared to Col-0 (Fig 4C). These results indicate that RBOHF is important for resistance to *Pst* DC3000 bacteria and contributes more to pre-invasive than post-invasive defence responses.

### RBOHF is required for PAMP-triggered ROS production in guard cells

Surprisingly, while flg22 causes stomatal closure, ROS production in *rbohD* or *rbohF* guard cells upon flg22 activation has not been reported. Moreover, these two NADPH oxidases were not implicated in the ROS increase in guard cells after treatment with yeast elicitors (46). We analysed roGFP2-Orp1 fluorescence in guard cells of leaf discs after dual excitation at 405 nm and 488 nm (emission at 508-526 nm) by confocal laser scanning microscopy (Fig. S5A). In Col-0, roGFP2-Orp1 oxidation started to increase from 20 min after flg22 treatment (Fig. S5B). No significant difference in roGFP2-Orp1 oxidation state was observed in *rbohD, rbohF, prx33-3* and *prx34-2* guard cells as compared to Col-0 in control conditions (Fig. 4D and Fig. S5C). The *rbohD* mutant was not impaired in flg22-mediated roGFP2-Orp1 oxidation in guard cells (Fig. 4D). By contrast, the *rbohF* mutant exhibited a marked increase in roGFP2-Orp1 oxidation compared to Col-0 at 30 and 60 min after flg22 treatment. Similarly, the *prx33-3* and *prx34-2* mutants which are impaired in stomatal immunity (11) showed a significant increase in flg22-induced roGFP2-Orp1 oxidation in guard cells as compared to Col-0 (Fig. S5C). These unexpected results suggest that a defect in PAMP-mediated stomatal closure is correlated with an enhanced H_2_O_2_ or oxidation state of guard cell cytosol/nucleus. A more detailed comparison of roGFP2-Orp1 oxidation and stomatal aperture of individual guard cells in Col-0 WT after control or flg22 treatment for 30 and 60 min shows that while roGFP-Orp1 is relatively reduced in control condition (Fig. 4E), there was no correlation between stomatal aperture and roGFP2-Orp1 oxidation (R = 0.0651, *P* = 0.4833). We found a negative correlation, albeit weak, between stomatal aperture and roGFP2-Orp1 oxidation across all flg22 treatments (flg22 30 min, R = −0.2428, *P* = 0.01492 and flg22 60 min, R = −0.2395, *P* = 0.01062) (Fig. 4E). As a complementary approach, we used H_2_DCFDA to analyse PAMP-induced ROS production in guard cells of NADPH oxidase mutants. H_2_DCFDA assays indicate that although the basal level of ROS was decreased in both *rbohD* and *rbohF* mutants, *rbohF* has a significantly smaller increase in ROS compared to Col-0 by 30 min after flg22 treatment (Fig. 4F). These results suggest that only RBOHF contributes to flg22-mediated ROS production in guard cells but RBOHD does not. Importantly, based on H_2_DCFDA assays, PRX33 and PRX34 are also implicated in PAMP-mediated ROS accumulation in guard cells (11). Therefore, across these mutants, ROS measurement by H_2_DCFDA shows a correlation between defects in ROS production and defects in flg22-induced stomatal closure while, particularly in the case of *rbohF*, cytosolic roGFP2-Orp1 is more oxidised. We also tested the stomatal response of *rbohD* and *rbohF* mutants to H_2_O_2_ which closes stomata through the activation of Ca^2+^ channels and inhibition of the inward K^+^ channels (49, 50). The results indicate that *rbohF* was partially impaired in H_2_O_2_ -mediated stomatal closure while WT stomatal closure was observed for the *rbohD* mutant (Fig 4G). Therefore, RBOHF may also act downstream of H_2_O_2_ during PAMP-mediated stomatal closure. As a control, we confirmed that *rbohF*, but not *rbohD*, is partially impaired in abscisic acid-mediated stomatal closure (Fig. S6A; (45, 48).

Moreover, as previously observed (46), these NADPH-oxidases are not involved in salicylic acid-mediated stomatal closure (Fig. S6B).

## Discussion

### Unlike BAK1 and BIK1, NADPH oxidases and PRXs are not fully required for PAMP-induced cytosolic hydrogen peroxide production

It was previously noted that flg22 caused a rapid luminol-measured burst followed by delayed oxidation of cytosolic roGFP2-Orp1 but the relationship between these events was not resolved (18). The use of mutants in apoplastic ROS production and protein kinases BIK1 and BAK1 involved in PAMP perception has provided additional information. As expected, the flg22-induced apoplastic ROS burst measured by luminol was rapid and transient and strictly dependent on RBOHD but not RBOHF. Surprisingly, the subsequent oxidation of cytosolic roGFP2-Orp1 was independent of both isoforms but notably, in the *rbohD* mutant, there was a delay in cytosolic roGFP2-Orp1 oxidation, suggesting that part of the cytosolic oxidation is dependent on H_2_O_2_ produced in the initial RBOHD-dependent burst. It is possible that other RBOH isoforms contribute to cytosolic roGFP2-Orp1 oxidation, but this is unlikely since DPI decreased the initial probe oxidation but did not affect it in the longer term. Therefore, it is apparent that PAMP treatment induces oxidation of roGFP2-Orp1 in an NADPH oxidase-independent manner. Activation of other apoplastic ROS producing enzymes, such as type III peroxidases is another possibility since they have been reported to be involved in the PAMP-induced ROS production (1, 2). Analysis of the four peroxidase mutants suggested that these isoforms do not play major roles in PAMP-induced roGFP2-Orp1 oxidation although PRX34 may contribute to sustain roGFP2-Orp1 oxidation in the long term and *PRX4* up-regulation boosted PAMP-triggered H_2_O_2_ production in the cytosol. Functional redundancy is likely to occur as PRXs belong to a large multigenic family of 73 members (51). Our results suggest that NADPH oxidases and apoplastic peroxidases may act additively but other enzymes play an important role in PAMP-mediated intracellular H_2_O_2_ signalling.

The cytosolic oxidation is pervasive because flg22 also oxidised GRX1-roGFP2, reaching a maximum slightly later. GRX1-roGFP2 is an indicator of glutathione redox state and this response may reflect increased use of glutathione in removing H_2_O_2_ through the various antioxidant systems (52, 53). In contrast to the *rboh* mutants, BAK1 was essential for PAMP-dependent roGFP2-Orp1 oxidation, while *bik1* was partially defective. The results raise questions about the source of H_2_O_2_ in the cytosol and how its production is activated downstream of these PTI regulators. The cytosol is not usually considered to produce significant amounts of ROS whilst the apoplast and organelles can. The treatments were carried out in the dark (in the plate reader), so photosynthesis is an unlikely source while mitochondrial electron transport and peroxisomal oxidases are established ROS sources (9).

However, PAMP treatment caused only a mild oxidation of mitochondrial roGFP2-Orp1 compared to cytosolic/nuclear roGFP2-Orp1, suggesting that mitochondria make a small contribution to cytosolic roGFP2-Orp1 oxidation (18). Further investigation is required. Conversely to activating H_2_O_2_ production, it is possible that BAK1/BIK1 inactivate H_2_O_2_ scavenging enzymes directly or indirectly, possibly *via* phosphorylation.

### *rbohF* is sensitive to oxidation caused by H_2_O_2_ and flg22

The results indicate that RBOHF is important for maintaining the function of the antioxidant system. Unlike *rbohD*, the *rbohF* mutant has a small increase in the oxidation state of cytosolic/nuclear roGFP2-Orp1 compared to Col-0, and importantly, when exposed to H_2_O_2_ in leaves and flg22 in guard cells, probe oxidation is greater than in Col-0. This conclusion is supported by decreased rosette size and increased bleaching of older leaves seen in *rbohFcat2* double mutants (10), suggesting that *rbohF* mutant has a decreased capacity to remove the excess H_2_O_2_ produced by the *cat2* catalase mutant. Interestingly, the dichlorofluorescein ROS-sensitive dye used by Chaouch *et al*. (2012) was not able to detect differences in between Col-0 and *rbohF* as compared to our measurements with roGFP2-Orp1. These results are consistent with a less active antioxidant system and, indeed, *rbohF* has lower expression of cytosolic ascorbate peroxidase (APX1), an enzyme known to be important in H_2_O_2_ removal (10). The small stature and stress sensitivity of *rbohF* suggests a role for ROS production by RBOHF in various aspects of plant growth and development (54, 55) and control over the antioxidant system. The higher pH of the apoplast of *rbohF* compared to Col-0 could be a factor in reduced growth (56).

### The role of RBOHF in stomatal defences

PAMP-mediated stomatal closure is considered to involve ROS production (57). Previous reports suggest that the RBOHD isoform is required for closure, presumably mediated by ROS production initially in the apoplast (7, 8, 44, 45). However, the defect in PAMP-mediated stomatal closure observed in *rbohD* mutant was not correlated with increased susceptibility to *Pst* bacteria (Macho et al., 2012; Kadota et al., 2014). A more recent study showed that RBOHD and RBOHF are not involved in flg22-triggered stomatal closure (58). Contrary to previous work, and by using different mutant alleles, we found that stomatal closure in response to flg22 and to *Pst* COR^-^ bacteria depends on RBOHF but not on RBOHD. Correspondingly, there was increased bacterial growth in *rbohF* mutant but not *rbohD*. It should be noted that the *rbohD* and *rbohF* mutants showed the expected responses for the apoplastic burst in response to flg22. Our results and previous data suggest that the relative contribution of RBOHD and RBOHF in stomatal immunity likely depend on experimental conditions. While unquantified differences in growth conditions may be involved, it should also be noted that immune responses are strongly dependent on the circadian clock that could for example mean that different NADPH oxidase isoforms to influence responses at different times of day (59).

The relationship between ROS production and stomatal closure was complex. The widely used probe H_2_DCFDA showed decreased ROS production in *rbohF* guard cells in response to flg22, which is commensurate with defect in stomatal closure. roGFP2-Orp1 and GRX1-roGFP2 have been used to demonstrate oxidation in guard cell cytosol during NaHS-induced stomatal closure (21). However, during the flg22 response, roGFP2-Orp1 oxidation state did not show a strong relationship with aperture and the probe was more oxidised in *rbohF* guard cells despite a lower H_2_DCFDA oxidation. Added to this, stomatal aperture in Col-0 had a poor correlation with roGFP2-Orp1 oxidation state in control condition, a phenomenon somewhat previously observed. Using epidermal strips incubated in stomatal opening buffer, the stomatal aperture seems not affected by high roGFP2-Orp1 oxidation in guard cells (21). A possible explanation is that roGFP2-Orp1 is specifically reporting cytosolic H_2_O_2_ which could be derived from multiple compartments. The use of catalase mutants has shown that, despite higher intracellular H_2_O_2_, stomatal aperture is unaffected although still responsive to exogenous H_2_O_2_ (60). This may explain why stomata remain open despite photosynthetic ROS production in the light. Our results therefore support the uncoupling of cytosolic H_2_O_2_ from stomatal aperture and that the extracellular production of superoxide or H_2_O_2_ by RBOHF is critical for flg22-induced closure. The LRR receptor kinase HPCA1 is a proposed extracellular H_2_O_2_ sensor which, when oxidised on extracellular cysteines, activates calcium channels leading to increased intracellular Ca^2+^ (61). This sensor provides an explanation for the requirement of apoplastic H_2_O_2_ for stomatal closure. Intriguingly, our results suggest that the ROS probe H_2_DCFDA could indicate that ROS other than H_2_O_2_ are involved in triggering closure while roGFP2-Orp1 indicates cytosolic conditions that are at least partly independent of stomatal aperture. The results with guard cells therefore partly mirror the findings with whole leaves where, as discussed above, the RBOHD-dependent apoplastic oxidative burst measured by luminol is independent of subsequent prolonged oxidation of cytosolic roGFP2-Orp1.

### Putative role of RBOHF in PAMP-mediated apoplastic alkalinisation

The *rbohF* mutant was also affected in stomatal closure induced by exogenous H_2_O_2_ suggesting that RBOHF function downstream or independently of ROS. Interestingly, despite a higher apoplastic pH in normal condition, the *rbohF* mutant was affected in flg22-induced apoplastic alkalinisation. In *rbohC* mutant, the amplitude of apoplastic pH fluctuations during root hair elongation is affected (Monshausen et al., 2007). It was shown recently that damage-induced membrane depolarization in roots is impaired in *rbohD* and *rbohF* mutants (Marhavý et al., 2019) and RBOHD might be important in propagating the systemic electric potential induced by wounding in leaves (Susuki et al, 2013). These results suggest that besides superoxide production, plant NADPH oxidases like their animal counterparts (38) are electrogenic through electron transport across the plasma membrane which is coupled to proton efflux probably through H^+^-ATPases (Fig S7). These extracellular H^+^ could participate in cell wall loosening *via* the expansin activities in normal conditions (56). During PAMP-triggered immunity, superoxide dismutation to H_2_O_2_, probably catalysed by unknown PAMP-activated superoxide dismutases or germin-like proteins, consumes protons and could increase apoplastic pH (Fig S7) which in turn activates apoplastic peroxidases (12). Alternatively, *rbohF* could have an indirect effect on the activity of plasma membrane H^+^- ATPases. It has been suggested that flg22 inhibits AHA1 and AHA2 H^+^- ATPase *via* dephosphorylation (62, 63). More importantly, apoplastic alkalinisation driven by RBOHF or AHA1/2 inhibition likely contributes to plasma membrane depolarisation which is well known to activates outward K^+^ channels and S-type anion channels leading to stomatal closure (64).

## Materials and Methods

### Plant materials and growth conditions

The roGFP2-Orp1 line (18), the GRX1-roGFP2 line (53) and all mutant lines are in *Arabidopsis* Col-0 background. *bik1* (SALK_005291), *rbohD* (CS9555), *rbohF* (CS9557), *prx4-2* (SALK_044730C), *prx33-3* (GK-014E05), *prx34-2* (GK-728F08) and *prx71-1* (SALK_123643C) were previously described (3, 11, 65). The *bak1-5* mutant was genotyped according to (25). The *rbohF* (SALK_034674) and *rbohD* (SALK_070610) lines were obtained from the European Arabidopsis Stock Centre. All T-DNA insertion mutants were confirmed by PCR genotyping prior to use (Supplemental Table 1). The genotyping of *rboh* mutants is shown in Fig. S8 and S9. F3 homozygous plants of the double transgenic lines made with roGFP2-Orp1 and the above mutants were generated by crossing homozygous parental lines. The progeny was selected on appropriate antibiotics and genotyping by PCR. Four- to five-week-old plants grown on soil in a growth chamber under short-day conditions (10 h light at 22°C/14 h dark at 19°C), at 60% humidity and illuminated with fluorescent tubes at 100 μmol m^−2^ s^−1^ light intensity were used for all the experiments.

### Statistical analysis

The experiments reported here were repeated at least three times with similar results unless otherwise mentioned. Non time course experiments were analysed by Student’s *t*-tests or ANOVAs followed by Tukey’s honestly significant difference (HSD) *post hoc* test using R software (R Core Team, https://www.R-project.org). The time course experiments, in which roGFP2-Orp1 and GRX1-roGFP oxidation state was followed after PAMP and H_2_O_2_ addition, were analysed by 2-way ANOVA using repeated measures for time. Significant differences between each treatment at each time were determined by Tukey’s multiple comparisons test using GraphPad Prism v8 (GraphPad, San Diego, California, USA). Raw data for stomatal aperture measurements are in Supplemental Table 2. All time course effects discussed in the text are significant (p<0.05) and the analysis is shown in Supplemental Table 3.

### Bacterial infection assay

The bacterial strain *Pst* DC3000 COR^-^ (DB4G3) (66) was cultivated overnight at 28°C in King’s B medium supplemented with Kanamycin and Rifampicin (each at 100 μg/mL). Bacteria were collected by centrifugation at 3000 *g* for 5 min at room temperature and washed twice in 10 mM MgCl2. Plants were surface-inoculated by spraying with a bacterial solution of 10^8^ cfu/ml in 10 mM MgCl2 containing 0.02% Silwet L-77, and plants were covered to maintain high humidity until disease symptoms developed. Alternatively, rosette leaves were syringe-infiltrated with a bacterial solution of 10^6^ cfu/ml in 10 mM MgCl2. After three days, bacterial growth in the apoplast of three leaves per plant and six plants per genotype was determined as previously described (67).

### Chemicals

Purified chemicals, except the flg22 and elf18 peptides (Peptron, Korea), were purchased from Sigma. Control solutions were stomatal buffer containing 1% ethanol for 2 mM salicylhydroxamic acid (SHAM) and 1 mM 2-Phenyl-4,4,5,5-tetramethylimidazoline-1-oxyl 3-oxide (cPTIO), 0.1% DMSO for 20 μM diphenyleneiodonium chloride (DPI), and water for 1 μM sodium azide, 10 μM to 100 mM hydrogen peroxide (H_2_O_2_), 50 mM 1,4-Dithiothreitol (DTT), 50 μM to 50 mM sodium nitroprusside (SNP), 10 nM to 10 μM flg22 or elf18.

### Measurements of stomatal aperture

Stomatal experiments were conducted as previously described (42). Epidermal peels collected from the abaxial side of young fully expanded leaves or leaf discs were floated in stomatal buffer (10 mM MES-KOH pH 6.15, 30 mM KCl) for 2.5 h under light (100 μmol m^−2^ s^−1^) to ensure that most stomata were opened before treatments. Solutions of chemicals or bacterial suspension at 10^8^ cfu.ml^−1^ in 10 mM MgCl2 were directly added in the stomatal buffer. After treatments, epidermal peels or leaf discs were further incubated under light for 2 h and observed under a light microscope (Carl Zeiss, Axioplan 2). Stomatal apertures of stomata in random areas were measured using ImageJ 1.42 software.

### Monitoring ROS in guard cells

2’,7’ dichlorofluorescein diacetate (H_2_DCFDA, Sigma) was used to measure ROS in guard cells (49). After 2.5 h incubation in stomatal buffer, epidermal peels were incubated with 50 μM H_2_DCFDA in 0.1% DMSO for 15 min. Excess H_2_DCFDA was then removed by washing 3 times for 20 min with stomatal buffer. Then, 5 μM flg22 or control solution were added. After 30 min incubation, H_2_DCFDA fluorescence was observed with a fluorescence microscope (Carl Zeiss, Axioplan 2) and fluorescence intensity of guard cells was analyzed using ImageJ software.

### Gene expression analysis in guard cell protoplasts

For each condition, about 50 young fully expanded leaves were immerged for 2 h in stomatal buffer containing 0.02% (v/v) Silwet-L77 with or without 1 μM flg22. Guard cell protoplasts were isolated as previously described (68) in the presence of transcriptional inhibitors 0.01% (w/v) cordycepin and 0.0033% (w/v) actinomycin D. For each condition, about 10^6^ guard cell protoplasts were obtained and their purity was above 98%. RNA was extracted using the RNeasy Plant Mini Kit with in-column Dnase 1 digestion (Qiagen). 200 ng of total RNA were reverse transcribed using 500 ng of oligo(dT)15 and the ImProm-II Reverse Transcription System following the manufacturer’s instructions (Promega). Quantitative real-time PCR reaction was performed on a Light Cycler 2 (Roche) using 10 μL SYBR Premix Ex Taq (Takara), 2 μL of 2-fold diluted cDNA, and 0.5 μM of primers in a total volume of 20 μL per reaction. The cycling conditions were composed of an initial 20 s denaturation step at 95°C, followed by 45 cycles of 95°C for 7 s, 60°C for 10 s, 72°C for 13 s. A melting curve was run from 65°C to 95°C to ensure the specificity of the products. Data were analyzed with the delta Ct method. Ubiquitin 1 (*UBQ1*) was used as a reference gene for normalization of gene expression levels. Control treatment was considered as expression level=1. As controls, the expression of the reference gene *ACT2* did not change between control and flg22 conditions while the expression of the PTI marker gene *PRX4* was induced by flg22 (Fig S4D). qRT-PCR primer sequences are listed in Supplemental Table 1.

### Multiwell plate reader-based fluorimetry

Because of random silencing of the transgene after 3 weeks of growth, plants expressing roGFP2-Orp1 were first selected with a epifluorescence binocular microscope. Leaf discs (6 mm diameter) were placed in a 96-well plate, immersed in 200 μl 10 mM MES-KOH pH 6.15, 30 mM KCl with their abaxial side facing up, and incubated for 2 h at 21 °C under laboratory lighting (PPFD ~10 μmol m^−2^ s^−1^) for recovery after wounding. roGFP2-Orp1 was excited sequentially at 400 nm ± 8 nm and 485 nm ± 8 nm in a CLARIOstar plate reader (BMG Labtech) and emission was recorded at 525 nm ± 20 nm with a gain set at 2000 and 1500 for the 400 and 485 nm excitations respectively. Each leaf discs were scanned from the top with the fluorescence recorded and averaged from 76 flashes per well organised as a spiral of 5 mm diameter. The initial 400/485 ratio of the resting state of leaf discs was estimated by reading the wells for 15 min before treatment. For each treatment, the emission of six Col-0 (WT) leaf discs was averaged and subtracted for all the data points to correct for background fluorescence. Changes in fluorescence over time were expressed relative to the initial ratio Ri as R/Ri to allow for differences in resting 400/485 ratio between leaf discs

### Luminol assay

Leaf discs of 6 mm diameter were cut in 4 equal pieces, immersed in water and incubated for 3 h minimum at room temperature for recovery after wounding. Before starting the assay, water was exchanged by a solution containing 100 μM luminol and 10 μg/mL horseradish peroxidase (≥ 250 units/mg solid). After adding control solution or 1 μM flg22, the luminescence was measured immediately using a CLARIOstar plate reader (BMG Labtech) with a reading time of 2 sec.

### Confocal imaging and image analysis

Leaf discs from rosette leaves were immersed in 10 mM MES-KOH pH 6.15, 30 mM KCl, incubated for 2 h at 21 °C under laboratory lighting (PPFD ~10 μmol m^-2^ s^-1^) for recovery after wounding and subsequently treated with control solution or 1 μM flg22 for 30 and 60 min. Leaf discs were mounted under a Zeiss confocal microscope LSM 880. Images were collected with a 20X lens (Plan-Apochromat, 0.8 numerical aperture) and roGFP2-Orp1 was excited sequentially at 405 and 488 nm and emission was detected at 508-526 nm, with the pinhole set to 1 airy unit. Single plane images were processed with ImageJ software. Background fluorescence was insignificant and not subtracted. Fluorescence intensity values of individual guard cells selected as region of interest was quantified on 32-bit converted images and the ratio 405/488 nm was calculated.

### Ratiometric pH quantification using Oregon green

Fully expanded leaves were syringe-infiltrated with 25 μM of Oregon Green 488 dextran (ThermoFisher) (39, 40). For the autofluorescence background, milliQ water was infiltrated into the apoplast. Plant were kept in the growth chamber for 2 h until excess water has evaporated. Oregon Green-treated leaf discs were placed in a 96-well plate, immersed in milliQ water and further incubated for 2 h at 21 °C under laboratory lighting (PPFD ~10 μmol m^−2^ s^−1^) for recovery after wounding. Oregon Green was excited sequentially at 440 nm ± 8 nm and 495 nm ± 8 nm in a CLARIOstar plate reader (BMG Labtech) and emission was recorded at 525 nm ± 20 nm with a gain set at 1250 and 1000 for the 440 and 495 nm excitations respectively. Each leaf discs were scanned from the top with the fluorescence recorded and averaged from 76 flashes per well organised as a spiral of 5 mm diameter. The initial 495/440 ratio of the resting state of leaf discs was estimated by reading the wells for 15 min before treatment. The emission of water-infiltrated leaf discs was averaged and subtracted for all the data points to correct for background fluorescence. To convert the fluorescence 495/440 ratio into pH values, a calibration curve with pH-buffered Oregon Green solution infiltrated into the apoplast of leaves was performed according to (McLachlan et al., 2016). The Boltzmann fit was chosen for fitting sigmoidal curves to calibration data and pH was determined according to the equation:

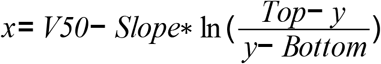

where x = pH, y = 495/440 ratio, Bottom = 0.4833, Top = 2.045, V50 = 4.513 and Slope = 0.6267. Changes in pH over time were normalised to the initial pHi as pH/pHi.

## Acknowledgements

The Biotechnology and Biological Sciences Research Council (BBSRC) provided funding (BB/I020004/1 and BB/N001311/1). We thank Murray Grant (University of Warwick, UK) for providing the *bik1* and *bak1-5* mutants and Markus Schwarzländer (University of Münster, Germany) for the kind gift of the roGFP2-Orp1 line.

---

**Table S1.** PCR primer sequences

**Table S2.** Stomatal aperture raw data.

**Table S3.** Statistical analysis of time course data.

**Fig. S1.** elf18-triggered roGFP2-Orp1 oxidation in mutants of PTI regulators, NADPH oxidases and apoplastic peroxidases.

**Fig. S2.** *In vivo* characterisation of roGFP2-Orp1 and GRX1-roGFP2 responses to the nitric oxide donor SNP.

**Fig. S3.** The response of roGFP2-Orp1 to the PAMP flg22 is not affected by the NO scavenger cPTIO.

**Fig. S4.** The NADPH oxidase RBOHF activates stomatal immunity.

**Fig. S5.** flg22-induced roGFP2-Orp1 oxidation in guard cells.

**Fig. S6.** The *rbohF* mutant is partly defective in ABA-mediated stomatal closure.

**Fig. S7.** Hypothetical model of the regulation of apoplastic pH by RBOHF during PTI activation.

