## Supplemental Figures 1-9 for "Differences between apoplastic and cytosolic reactive oxygen species production in *Arabidopsis* during pattern-triggered immunity"

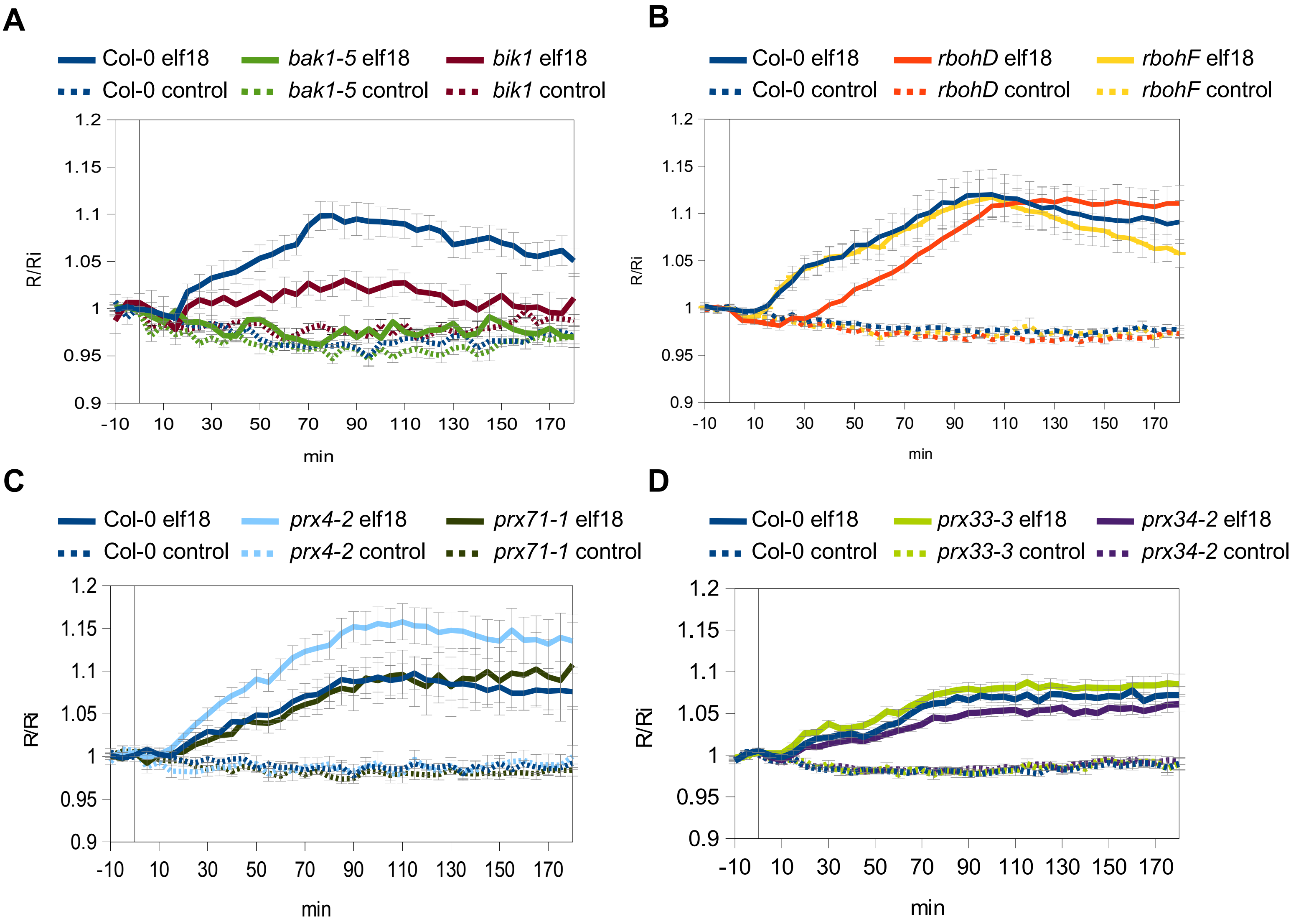


**Fig. S1.** elf18-triggered roGFP2-Orp1 oxidation in mutants of PTI regulators, NADPH oxidases and apoplastic peroxidases

(A-D) Kinetics of roGFP2-Orp1 oxidation in leaves of *bak1-5* and *bik1* (A), *rbohD* and *rbohF* (B), *prx33-3* and *prx34-2* (C), and *prx4-2* and *prx71-1* (D) mutants in response to elf18. Leaf discs were exposed at t = 0 min to control solution or 1 µM elf18. The 400/485 nm fluorescence ratio (R) was measured over time by multiwell fluorimetry (excitation at 400 ± 8 and 485 ± 8 nm; emission, 525 ± 20 nm) and expressed relative to the mean initial ratio (Ri) before treatment (R/Ri). Data are means ± SE from at least three independent experiments (n ≥ 16, B-D). In (A), a representative experiment is shown (n = 6).

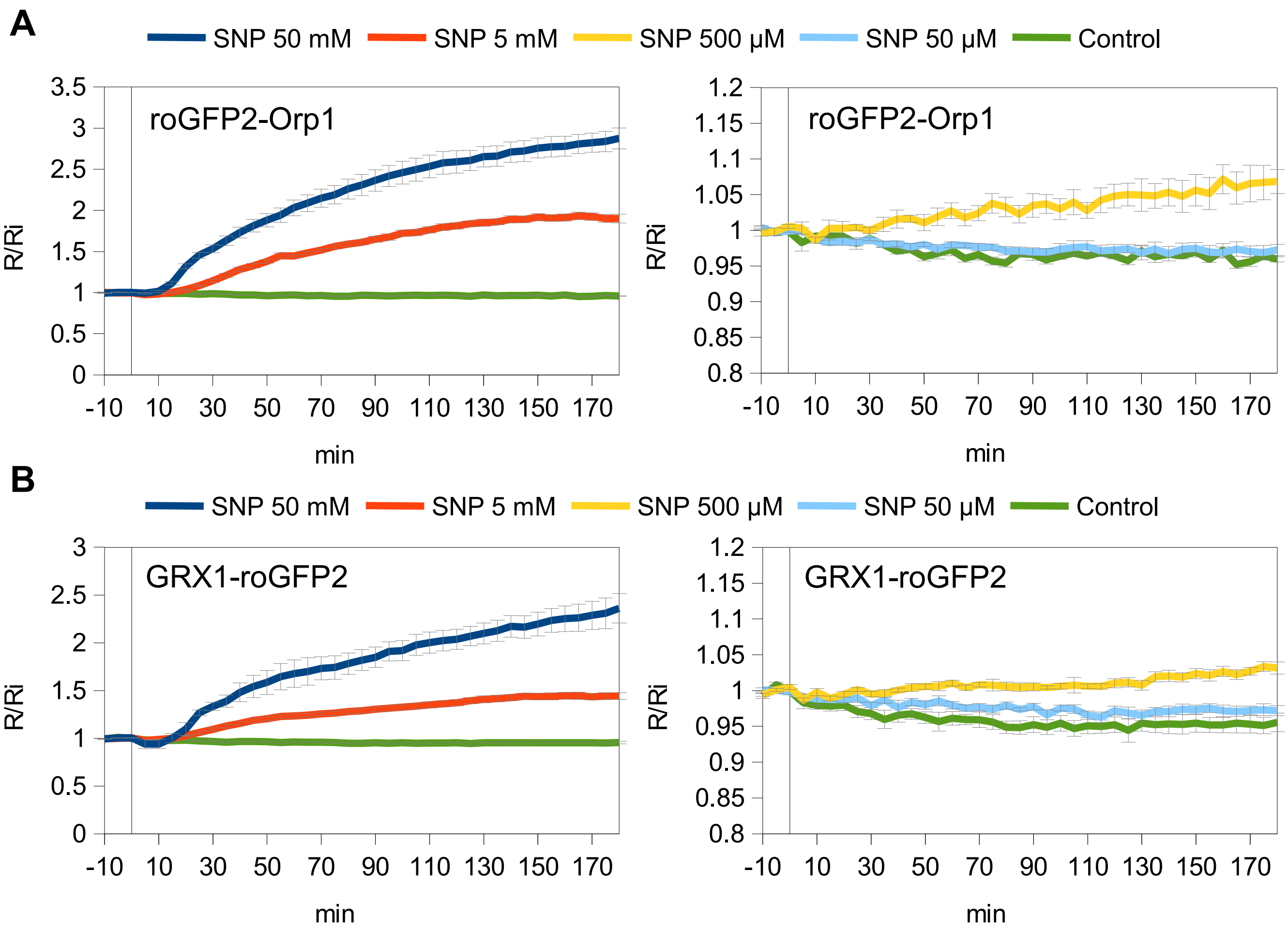


**Fig. S2.** *In vivo* characterisation of roGFP2-Orp1 and GRX1-roGFP2 responses to the nitric oxide donor SNP.

(A-B) Kinetics of roGFP2-Orp1 (A) and GRX1-roGFP2 (B) oxidation in leaves in response to NO. Leaf discs were exposed at t = 0 min to control solution or various concentration of the NO donor SNP. The 400/485 nm fluorescence ratio (R) was measured over time by multiwell fluorimetry and expressed relative to the mean initial ratio (Ri) before treatment (R/Ri). For clarity high concentration of SNP are presented on the left panels and lower doses are presented on the right panels. Data are means ± SE from a representative experiment (n = 6). The experiments have been repeated at least twice with similar results.

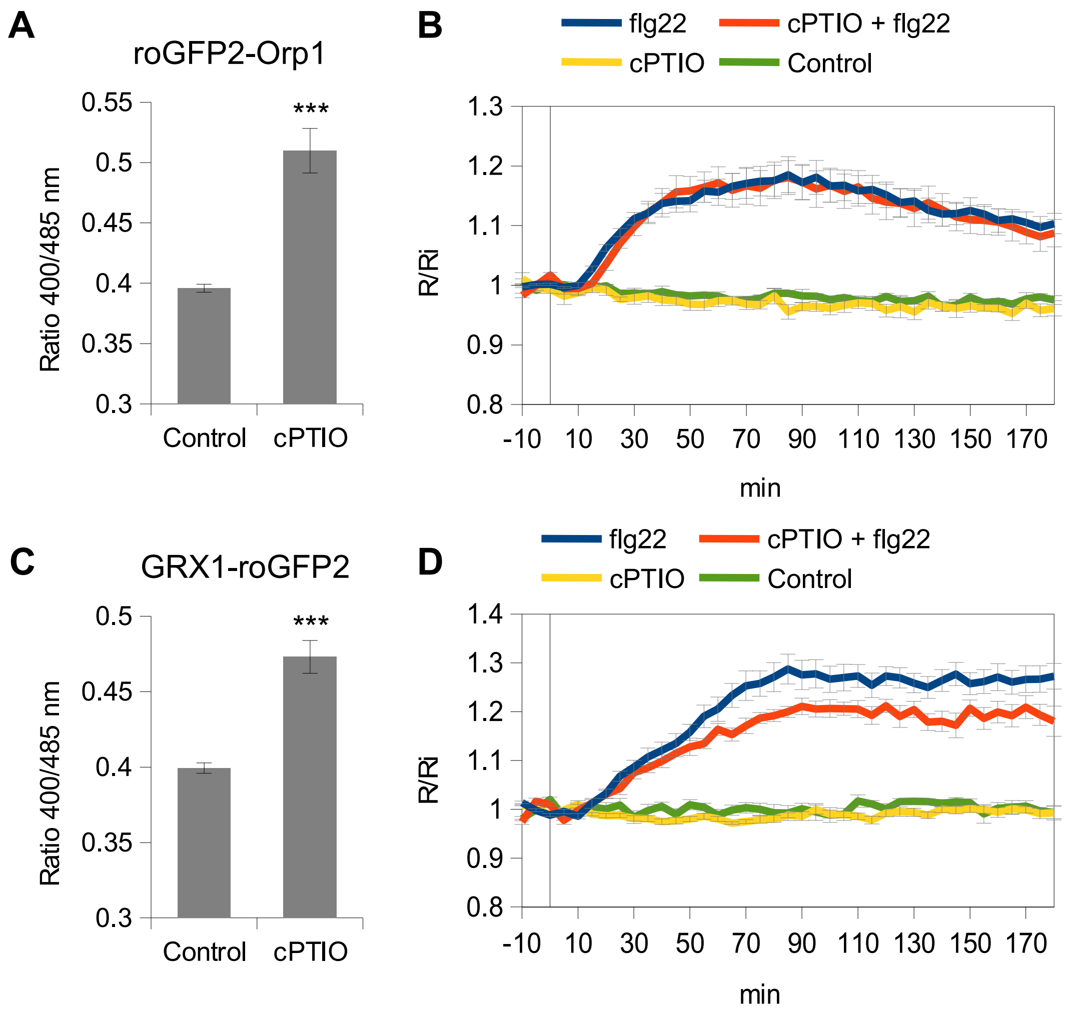


**Fig. S3.** The response of roGFP2-Orp1 to the PAMP flg22 is not affected by the NO scavenger cPTIO. (A-C) Effect of the NO scavenger cPTIO on the oxidation state of roGFP2-Orp1 (A) and GRX1-roGFP2 (C). Leaf discs were treated for 2 hrs with 1 % ethanol as a Control or 1 mM cPTIO and the oxidation status of roGFP2-Orp1 and GRX1-roGFP2 (ratio 400/485 nm) was measured by multiwell fluorimetry. Data are means ± SE from a representative experiment (n = 12). Asterisks indicate statistically significant differences between Control and cPTIO treatments based on a two-tailed Student's t-test (***$P$<0.001).

(B-D) Effect of cPTIO on flg22-induced oxidation of roGFP2-Orp1 (B) and GRX1-roGFP2 (D). After 2 hrs of pre-treatment with Control solution or 1 mM cPTIO, leaf discs were exposed at t = 0 min to Control solution or 1 µM flg22. The 400/485 nm fluorescence ratio (R) was measured over time by multiwell fluorimetry and expressed relative to the mean initial ratio (Ri) before flg22 treatment. Data are means ± SE from a representative experiment (n = 6).

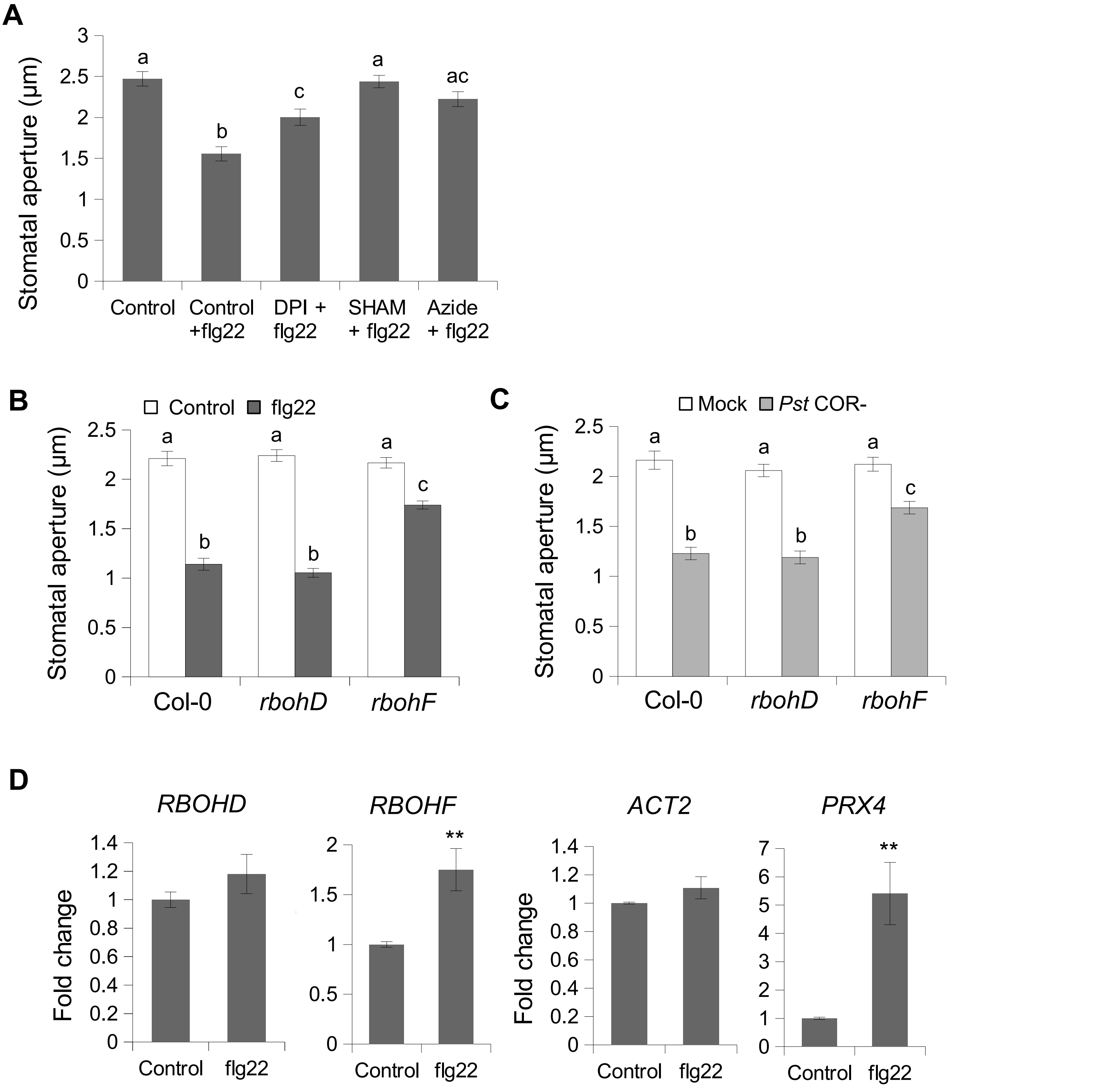


**Fig. S4.** The NADPH oxidase RBOHF activates stomatal immunity
(A) Stomatal aperture in epidermal peels of Col-0 WT pre-treated for 30 min with Control (0.1% DMSO), 20 μM DPI, 2 mM SHAM, or 1 μM sodium azide and exposed to Control solution or 5 μM flg22 for 2 h. Data are means ± SE (n ≥ 80) from a representative experiment.
(B) Stomatal apertures in WT Col-0, *rbohD*, and *rbohF* epidermal peels exposed to Control solution or 5 μM flg22 for 2 h.
(C) Stomatal apertures in Col-0 WT, *rbohD*, and *rbohF* epidermal peels exposed to Mock control (10 mM $MgCl_2$) or $10^8$ cfu/ml COR-deficient *Pst* DC3000 (*Pst* $COR^-$) bacteria for 2 h. In (B) and (C) data are means ± SE (n ≥ 100) from a representative experiment. Different letters indicate significant differences at $P < 0.001$ (B-C) and $P < 0.05$ (A) based on a Tukey's HSD test.
(D) *RBOHF* expression is induced by flg22 in guard cells. Expression analysis of *RBOHD*, *RBOHF,* the reference gene *ACT2,* and the PTI marker gene *PRX4* by RT-qPCR in guard cell protoplasts isolated from leaves of 5-week-old Col-0 plants incubated in stomatal buffer for 2 h without (Control) or with 1 μM flg22. Transcript levels were normalized to *UBQ1*. The changes in transcript levels are relative to Control treatment (expression value = 1). Error bars indicate SE of six independent experiments (n = 6). Asterisks indicate statistically significant differences between Control and flg22 treatments based on a two-tailed Student's t-test (**P<0.01).

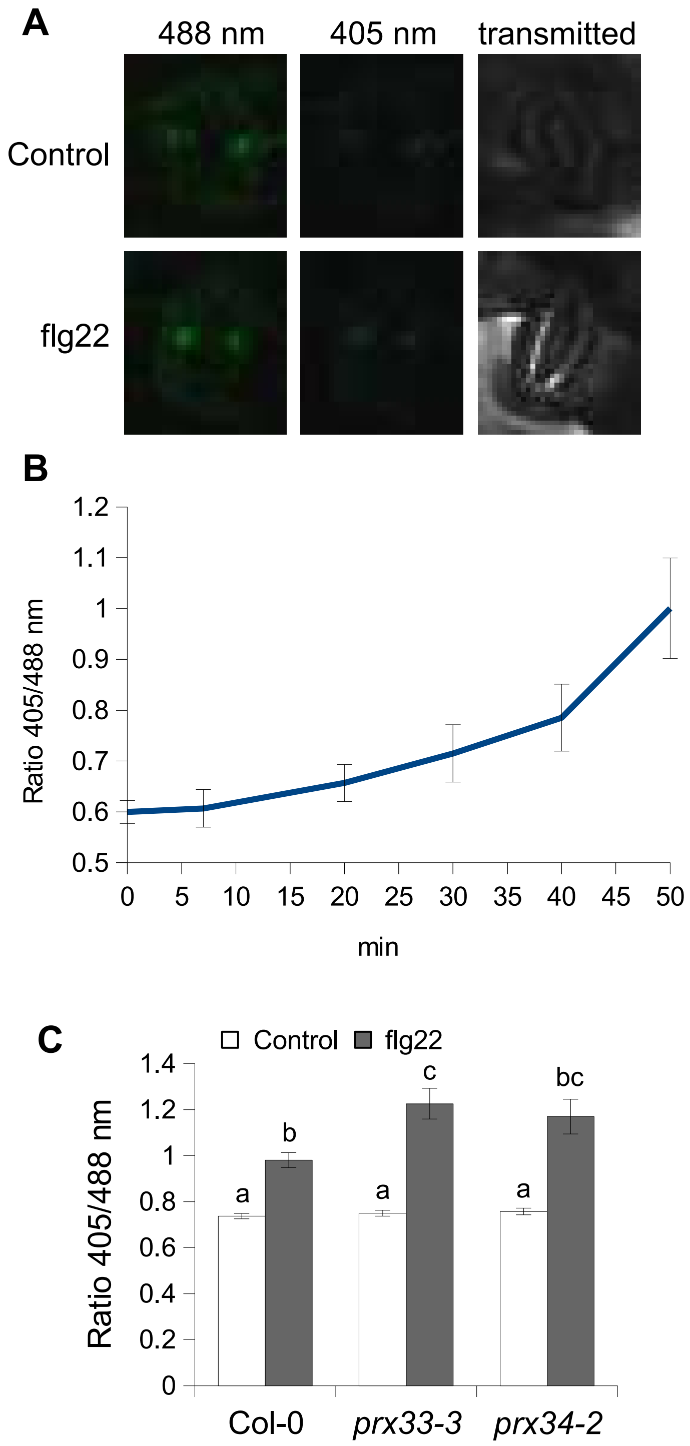


**Fig. S5.** flg22-induced roGFP2-Orp1 oxidation in guard cells
(A) Representative images of the fluorescence emission at 517 ± 9 nm following excitation at 488 and 405 nm in selected Col-0 WT guard cells after treatment with Control solution or 1 µM flg22 for 1 h.
(B) Kinetics of roGFP2-Orp1 oxidation in WT guard cells in response to flg22. Leaf discs from Col-0 WT were exposed at t = 0 min to 1 µM flg22 and the ratio 405/488 nm of guard cells was measured over time by confocal microscopy. Data are means ± SE from a representative experiment (n = 7).
(C) roGFP2-Orp1 oxidation state in guard cells of Col-0 WT, *prx33-3* and *prx34-2* mutants after flg22 treatment. Leaf discs were exposed to Control solution or 1 µM flg22 for 60 min. The ratio 400/488 nm of stomata selected as ROIs was quantified from images of the fluorescence emission at 517 ± 9 nm following excitation at 488 and 405 nm. Data are means ± SE (n ≥ 60 guard cells) from a representative experiment. Different letters indicate significant differences at $P < 0.01$ based on a Tukey's HSD test.

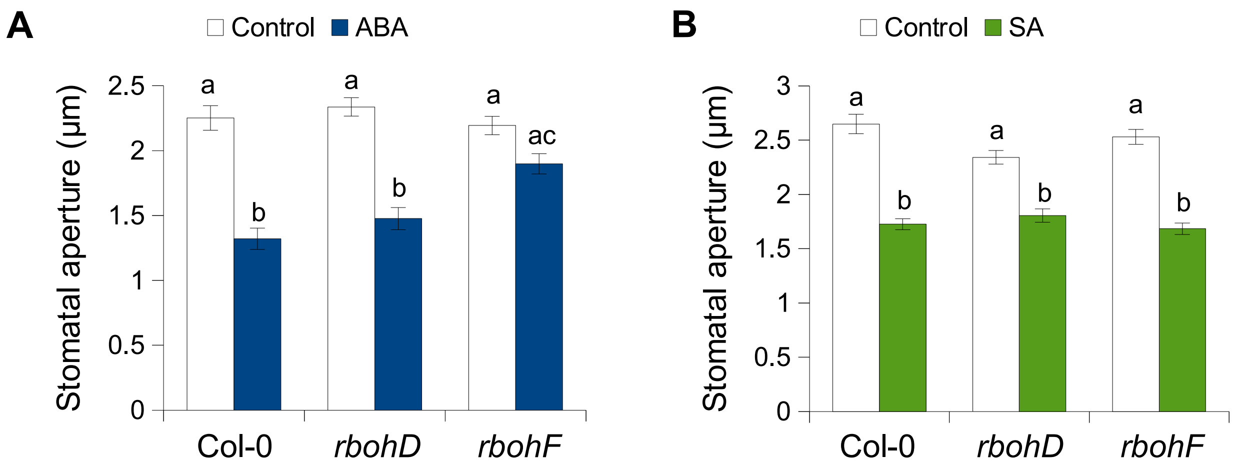


**Fig. S6.** The *rbohF* mutant is partly defective in ABA-mediated stomatal closure

(A) Stomatal apertures in Col-0 WT, *rbohD*, and *rbohF* epidermal peels exposed to Control solution (0.01% ethanol) or 1 μM abscisic acid (ABA) for 2 h.

(B) Stomatal apertures in WT Col-0, *rbohD*, and *rbohF* epidermal peels exposed to Control solution (0.01% ethanol) or 10 μM salicylic acid (SA) for 2 h. In (A) and (B) data are means ± SE ($n \geq 100$) from a representative experiment and different letters indicate significant differences at $P < 0.01$ (A) and $P < 0.001$ (B) based on a Tukey's HSD test.

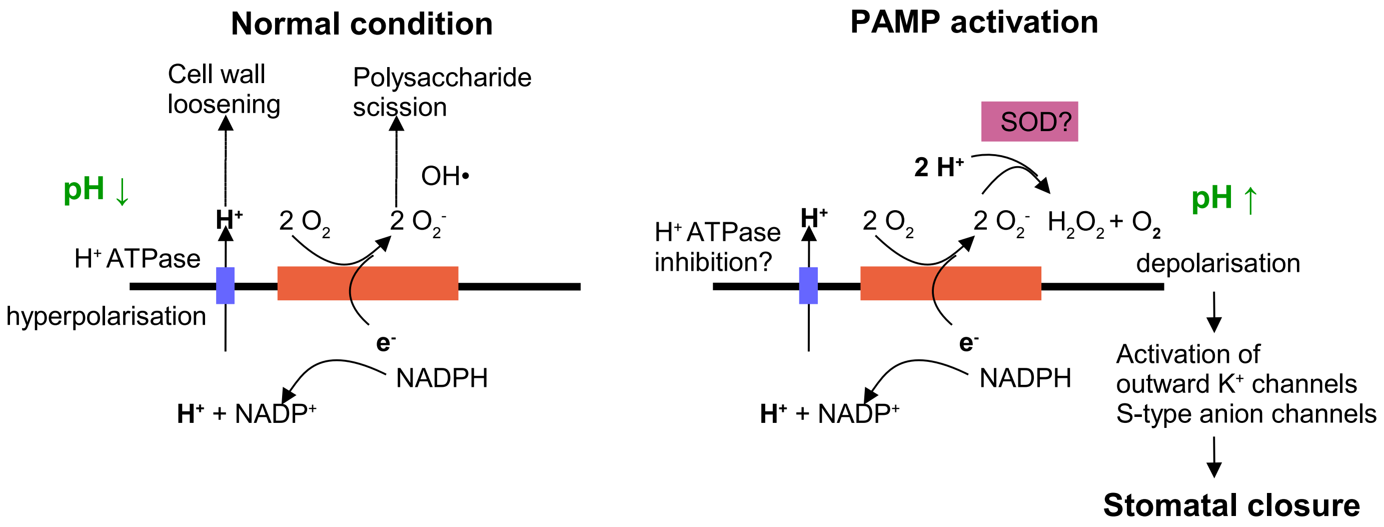


**Fig. S7.** Hypothetical model of the regulation of apoplastic pH by RBOHF during PTI activation. In normal unstressed condition, the efflux of electron ($e^-$) produced by the NADPH oxidase activity is compensated by an efflux of proton ($H^+$) probably generated by plasma membrane $H^+$ATPases. Both the acidification of the apoplast and the production of hydroxyl radicals (OH•) from superoxide ($O^{2-}$) through the Haber–Weiss and Fenton reactions contribute to cell expansion. The plant immune response may induces the inhibition of $H^+$ATPases and the activation of RBOHF together with superoxide dismutases (SOD) or germin-like proteins which dismutates $O^{2-}$ to $H_2O_2$, a reaction that consumme $H^+$. The resulting alkalinisation of the apoplast induces a depolarisation of the plama membrane ultimately leading to stomatal closure via the activation of ion channels.

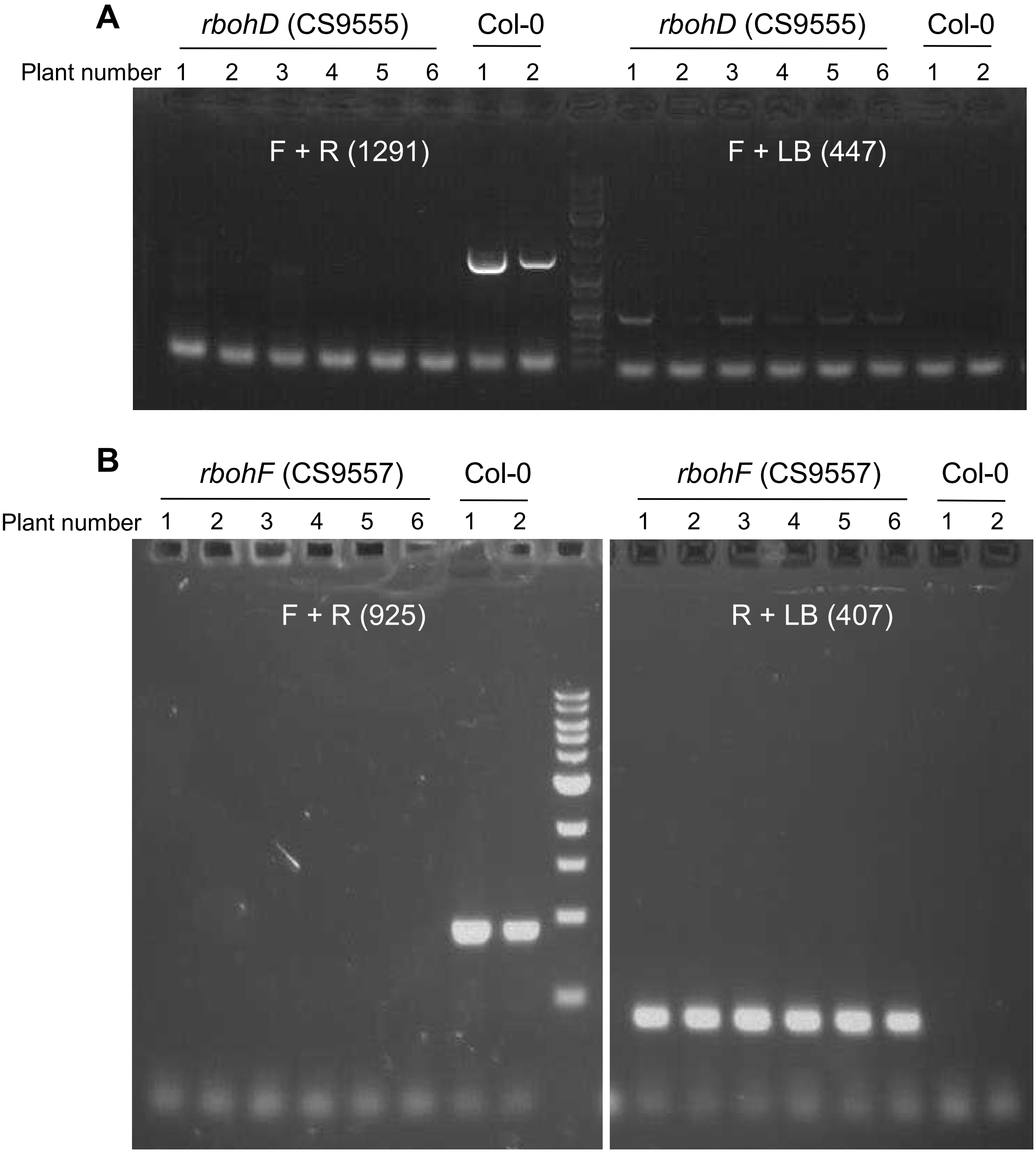


**Fig. S8.** Genotyping by PCR of *rbohD* and *rbohF dSpm* transposon mutants
(A-B) Representative images of PCR for genotyping the *rbohD* CS9555 (A) and *rbohF* CS9557 (B) mutants (Torres et al., 2002). Wild-type allele was amplified with gene specific forward and reverse primers (F + R), and T-DNA mutant allele was amplified with a gene specific primer (F or R) and the DsLox T-DNA left border (LB) primer. The expected size of amplicons is indicated under brackets. Col-0 wild-type genomic DNA was used as a control.

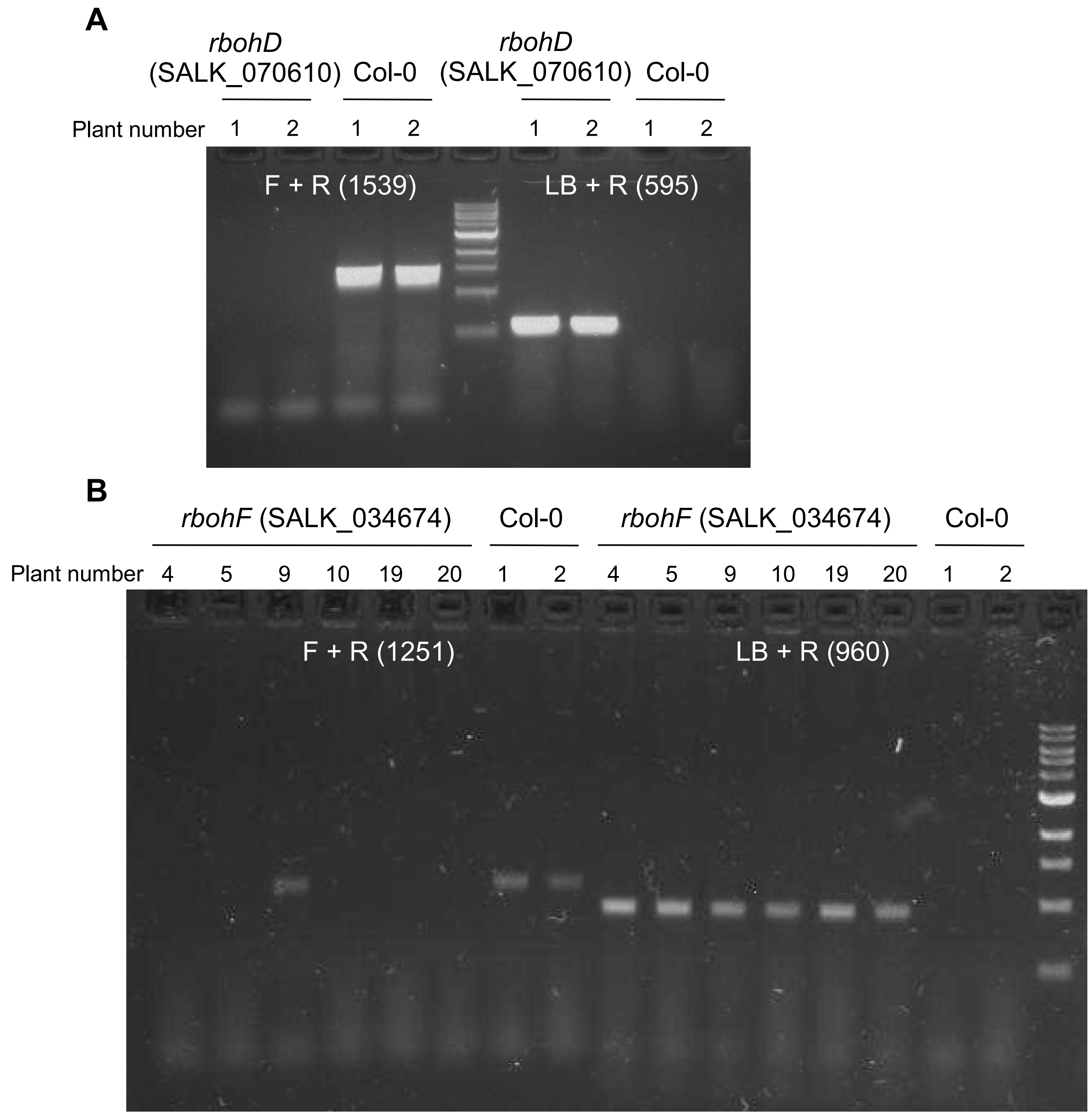


**Fig. S9.** Genotyping by PCR of *rbohD* and *rbohF* Salk T-DNA mutants
(A-B) Representative images of PCR for genotyping the *rbohD* SALK_070610 (A) and *rbohF* SALK_034674 (B) mutants. Wild-type allele was amplified with gene specific forward and reverse primers (F + R), and T-DNA mutant allele was amplified with a gene specific reverse primer (R) and the Salk T-DNA left border (LB) primer. The expected size of amplicons is indicated under brackets. Col-0 wild-type genomic DNA was used as a control.
